# Machine learnt image processing to predict weight and size of rice kernels

**DOI:** 10.1101/743427

**Authors:** Samrendra K Singh, Sriram K Vidyarthi, Rakhee Tiwari

**Affiliations:** CNH Industrial, Burr Ridge, IL 60527, USA; The Morning Star Company, Woodland, CA, 95695, USA; Department of Biological and Agricultural Engineering, University of California, Davis, CA 95616, USA

**Author notes:** Corresponding author: Samrendra K Singh, Address: CNH Industrial, Burr Ridge, IL 60527, USA.

**Keywords:** Machine Learning, Rice, Image processing, Stacked ensemble model, Kernel Length

## Abstract

Accurate measurement of rice kernel sizes after milling is critical to design, develop and optimize rice milling operations. The size and mass of the individual rice kernels are important parameters typically associated with rice quality attributes, particularly head rice yield. In this study, we propose a novel methodology that combines image processing and machine learning (ML) ensemble to accurately measure the size and mass of several rice kernels simultaneously. We have developed an image processing algorithm with the help of recursive method to identify the individual rice kernels from an image and estimate the size of the kernels based on the pixels a kernel occupies. The number of pixels representing a rice kernel has been used as its digital fingerprint in order to predict its size and mass. We have employed a number of popular machine learning models to build a stacked ensemble model (SEM), which can predict the mass of the individual rice kernels based on the features derived from the pixels of the individual kernels in the image. The prediction accuracy and robustness of our image processing and SEM are quantified using uncertainty quantification. The uncertainty quantification showed 3.6%, 2.5%, and 2.4% for mean errors in estimating the kernel length of small-grain (Calhikari-202), medium-grain (Jupiter), and long-grain (CL153) rice, respectively. Similarly, mean errors associated with predicting the 1000 grain weight are 4.1%, 2.9%, and 4.3% for Calhikari-202, Jupiter, and CL153, respectively. Use of the developed algorithm in rice imaging analyzers could facilitate head rice yield quantifications and promote quicker rice quality appraisals.

## 1 Introduction

Rice (Oryza sativa) is a major agricultural commodity in the world. The total annual production of milled rice in the world and the US in 2018/19 was about 501 and 7.1 million metric tons, respectively (USDA FAS, 2019). Rice (or Paddy) is processed into a wide range of products, of which, milled rice is produced after removal of its outer husk (hull) and embryo during milling and is generally preferred for human consumption over other forms of rice in the most countries (Prakash and Pan, 2009). In this article, “rice” refers to milled rice, unless specified otherwise.

### 1.1 Importance and methods of determining the size and mass of milled rice kernels

Rice sizing is important which significantly affects the customer perception about rice classification and thus its pricing (Prakash, 2011; Singh et al., 2019). The rice kernel size is an important attribute that is used internationally to determine the class of the rice. Rice is generally classified in three categories based on kernel length or length/width ratio or combination of both - short kernel, medium kernel and long kernel. For example, according to one convention, a long kernel rice has a length-width ratio of 3.0 or more to 1, a medium kernel rice has 2 - 2.9 to 1 and a short kernel rice has 1.9 or less to 1 (FAO, 1995; USDA, 1994). The length, width and their ratio of rice kernels provide uniformity in the rice samples, which is an important aspect to customers.

Rice kernel size is also one of the major indicators of rice quality. Larger average kernel size may contribute to a higher milling yield (Whan, et al., 2014). Since rice is generally consumed as intact kernels, the importance of maintaining kernel size is even more important for the rice industry. Broken rice can be defined with lengths that are less than three-fourths of the unbroken kernel length, only has typically 60% to 80% of the market value of remaining rice that is known as head rice (Siebenmorgen et al., 2008).

Food sizes, including cereals, grains, fruits, vegetables and nuts are key factors in several food processing unit operations, such as drying, sorting, size reduction, canning, peeling, dicing, blanching and heat treatment operations (Vidyarthi et al., 2019). Size sorting of food is frequently used in food industry to accommodate various food processes. For instance, different tomato sizes can affect the heat transfer rate during peeling process, including steam, lye or infrared peeling and thus tomato size sorting is an important unit operation in industrial tomato peeling and canning processes (Li, 2012; Pan 2009; Vidyarthi, 2017; Vidyarthi et al., 2019), as well as, new food product development which is crucial to the sustainable growth of food industry (Vidyarthi and Evans, 2019; Xu et al., 2019) and sanitation and disinfection of fruits and vegetables (Deng et al., 2019).

Rice sizing is important which significantly affects the customer perception about rice classification and thus its pricing (Prakash, 2011; Singh et al., 2019). The rice kernel size is an important attribute that is used internationally to determine the class of the rice. Rice is generally classified in three categories based on kernel length or length/width ratio or combination of both -short kernel, medium kernel and long kernel. For example, according to one convention, a long kernel rice has a length-width ratio of 3.0 or more to 1, a medium kernel rice has 2 - 2.9 to 1 and a short kernel rice has 1.9 or less to 1 (FAO, 1995; USDA, 1994). The length, width and their ratio of rice kernels provide uniformity in the rice samples, which is an important aspect to customers.

Rice kernel size is also one of the major indicators of rice quality. Larger average kernel size may contribute to a higher milling yield (Whan, et al., 2014). Since rice is generally consumed as intact kernels, the importance of maintaining kernel size is even more important for the rice industry. Broken rice can be defined with lengths that are less than three-fourths of the unbroken kernel length, only has typically 60% to 80% of the market value of remaining rice that is known as head rice (Siebenmorgen et al., 2008).

Traditional measurement of rice kernel size has been tedious and slow due to the manual nature of measurements (e.g., use of calipers to measure kernels one at a time) and their scoring methods have been subjective in nature because they are mostly based on visual inspection (Ramya et al, 2010; Santos et al., 2018). Some commonly used measurement methods for determining seed size and mass, such as thousand-kernel weight or hectoliter weight are fast and not prone to error, however, they do not specify a variation within a sample. Yabe et al. (2018) used 128 Japanese rice varieties and assessed genetic improvement in kernel yield in rice by quantitatively analyzing the genotypic differences in kernel-filling ability and describing the kernel weight distribution, which is the probability density function of single kernel weight in a panicle. Another existing technology used for detailed characterization of seed shape is single kernel characterization system (SKCS); however, it has relatively low-throughput, and is a destructive method (Martin et al., 1993).

There is an increasing use of image analysis for kernel size measurement. Recent developments in imaging technologies have enabled kernel measurements fast with relatively higher throughput. Classification of the rice kernels can also be made with the help of image processing technologies. These technologies usually involve digital imaging of rice kernels, computer scanning and analysis of kernels using imaging software by calculating the kernel areas and pixel values. For example, SeedCount (Next Instruments, NSW, Australia) utilizes image analysis to measure the size of individual seeds in a sample, which allows to estimate the sample mean and variations accurately. However, this method is time consuming, especially for large number of samples and can also be cost prohibitive (Whan et al., 2014). ImageJ is general purpose image analysis software that has been used to analyze seed shape and size parameters in a range of kernels, including wheat and rice (Li et al., 2009). Smartgrain is another image analysis system establishes seed area, perimeter, width, and length to extract seed characteristics by building ellipses on identified kernels (Tanabata et al., 2012). SHAPE analyzes seed shape by producing elliptic Fourier descriptors of 2- and 3-dimensional characteristics from photographs of vertically and horizontally oriented seed; however, it is manual intensive and requires preparation of individual seeds (Williams et al., 2013). Whan et al. (2014) analyzed the measurements and color of wheat and *Brachypodium distachyon* seeds using GrainScan which utilized reflected light to capture color information from color space (CIELAB). However, all these existing methods of determining seed size are mostly in development stage and high-throughput measurement is still a challenge.

### 1.2 Machine learning in rice research

For the first time we employed machine learning (ML) using stacked ensemble model (SEM) in the field of rice processing. Machine learning models offer a unique way to create a predictive model system in the form a known data set (Bejagam et al., 2018). It has been successfully implemented by several researchers in the field of rice processing. Its application is primarily focused in the area of classifying the rice grains based on variety or milling quality (El-Telbany, 2006; Fayyazi et al., 2017; Gujjar and Siddappa, 2013; Guzman and Peralta, 2008; Kaur and Singh, 2013; Liu et al., 2005; Mousavi et al., 2012; Prajapati and Patel, 2013). Cheng and Matson (2015) used ML technique to differentiate between rice and weed. Zareiforoush et al. (2016) compared different ML methods for qualitative classification of the rice grains. Shiddiq et al. (2001) estimated the degree of rice milling using ML.

In order to evaluate the accuracy and robustness of the ML methods, we performed the uncertainty quantification (UQ) using bootstrapped sampling method. In this method, the original data set of size n is used to create N*_BS_* number of data sets, each of size n, using a technique known as bootstrapping. In bootstrapping, a new dataset is created by picking a data point randomly from the original data set. The process is repeated n times and a data point from the original data set may not be picked or picked more than once. A bootstrapped dataset may have a few duplicates. The errors or residuals distribution obtained with the N*_BS_* bootstrapped data set gives a more reliable indication regarding the errors and uncertainties associated with the model predictions.

Considering the importance of quantifying the dimensions and mass of the rice kernels, the current methods have time and consistency limitations. The traditional method of measuring the rice dimensions can be extremely tedious especially if done at large scales. We propose a novel and robust technique to accurately measure the dimensions along with the individual mass and other statistics of the rice kernels. Our methodology includes image processing of the rice kernel sample to identify the individual kernels and then use a ML based SEM to predict the mass of the individual kernels. The objectives of this paper are to demonstrate the application of image processing in combination with the SEM to accurately predict the size and mass of the rice kernels. Along with estimating the size and mass, we attempt to quantify the errors and uncertainty associated with the predictions.

## 2 Materials and methods

### 2.1 Rice samples

Based on the kernel size, milled rice of three popular US-grown rice cultivars (CL153, Jupiter, and Calhikari-202; all harvested in 2018) were selected for conducting the tests in this study. CL153, Jupiter, and Calhikari-202 are long-grain, medium-grain, and short-grain cultivars, respectively; these rice cultivars are grown in mid-South United States and California. The rice samples were procured in Ziploc^®^ bags from the department of Biological and Agricultural Engineering at the University of California, Davis which received the samples from the state of Arkansas, Louisiana and California in US, respectively, for research purposes.

### 2.2 Size and weight measurement of rice kernels

After procurement of rice samples, 55 rice kernels of each cultivar were spread on a flat dark grey surface divided into grids. We chose 55 kernels to ensure a balance between the resolution and overall number of images to be taken. A large number of kernels in a frame would result in lower resolution of the individual grains, whereas a lower count per image would lead to a greater number of images. Each kernel was placed into a separate grid. Five broken kernels were also selected and placed in the grids. The lengths and breadths of the kernels were measured individually using a Vernier caliper (Your Partner in Precision, Newton, MA; Model - SPI, 14-792-6) with a precision of 0.001 mm. Each kernel was weighed using a scale (Mettler Toledo; Model - ML503T) with a precision of 1 mg. This process was repeated 19 times for each cultivar to obtain measurements for 1005 rice kernels.

### 2.3 Image processing algorithm

The rice grains were placed on a dark grey surface to ensure a fair contrast between the background and the white kernels. A reference object (13 × 7 mm rectangular paper strip) was also placed in the same image, along with the rice kernel. A cellphone camera (Apple, Cupertino, CA; Model: iPhone 7 plus) was used to take an image of rice kernels, approximately at 0.2 m from the surface, with the camera flash turned off. The default image size from the camera was 3553 × 2013 pixels. The initial image from the camera was cropped to ensure that the image didn’t contain any objects other than the rice kernels and the reference paper strip. The cropped image was then converted into a grayscale image by taking average of r, g and b values of the pixels (Eq 1). Grayscaling the image allowed us to use a single threshold value to create a binary image. A binary image, as its name suggests, consists of either black (r = 0, g = 0, b = 0) or white pixels (r = 255, g = 255 and b = 255). The binary image is derived from the grayscale image by comparing every pixel with a threshold value (*Th_pixel_*). Any pixel with grayscale value below the threshold (*Th_pixel_*) was assumed to be background or black pixel and anything equal or above the threshold was assumed to be a pixel representing the rice kernel or white pixel (Eq 2). The choice of the *Th_pixel_* depends upon the lighting and contrast between the background and the rice kernel. The value of the *Th_pixel_* was adjusted manually for every image to cancel the background noise (due to dust, reflection etc) while maintaining the integrity of the rice kernels. This process creates standardized binary images of the rice kernels. The individual rice kernels from the image were then identified using recursive algorithm.

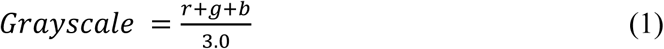

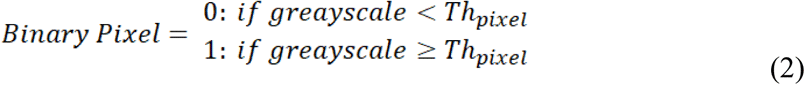

The recursive algorithm, available in most programing languages, enables a function to call itself from within (Arora et al, 2008). We designed the recursive function in such a way that it visits a pixel of white color and tags the previously untagged neighboring white-colored pixels by visiting these pixels as shown in Fig 1. While visiting each untagged neighboring pixel, it tags the untagged neighboring white-pixels again. This process continues until all the connected white-pixels are tagged. At the end of the function call, the algorithm counts the number of pixels tagged to identify a rice kernel in the image. This algorithm tags every white-pixel connected to each other. Irrespective of the starting pixel, the recursive method goes through every white-pixel in a cluster of connected pixels, where a cluster of white-pixels represents a single rice kernel in the binary image. The recursive function in this case ignores a previously tagged white-pixel. The algorithm scans the entire binary image until every white-pixel has been tagged. Each cluster at the end of the scanning is treated as a separate object.

**Fig. 1.**
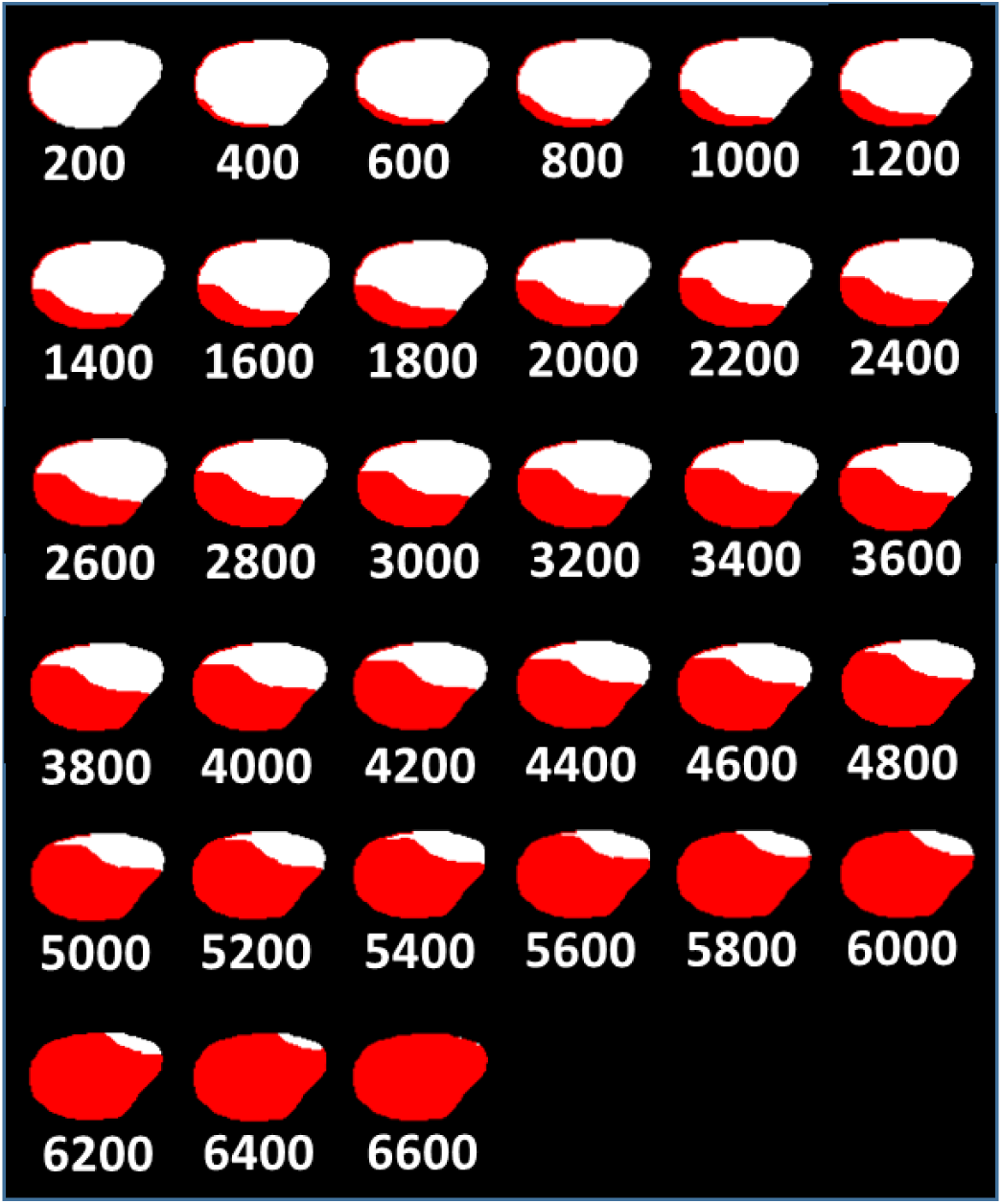
Step by step progression of recursive algorithm as it tags the white connected pixels of a rice kernel. The tagged pixels are recolored as red for clarity. The numbers below each image represents the number of recursive calls or number of white-pixels tagged. The recursive function stops when all connected white-pixels have been tagged. This example has 6600 pixels representing the rice kernel.

**Fig. 2.**
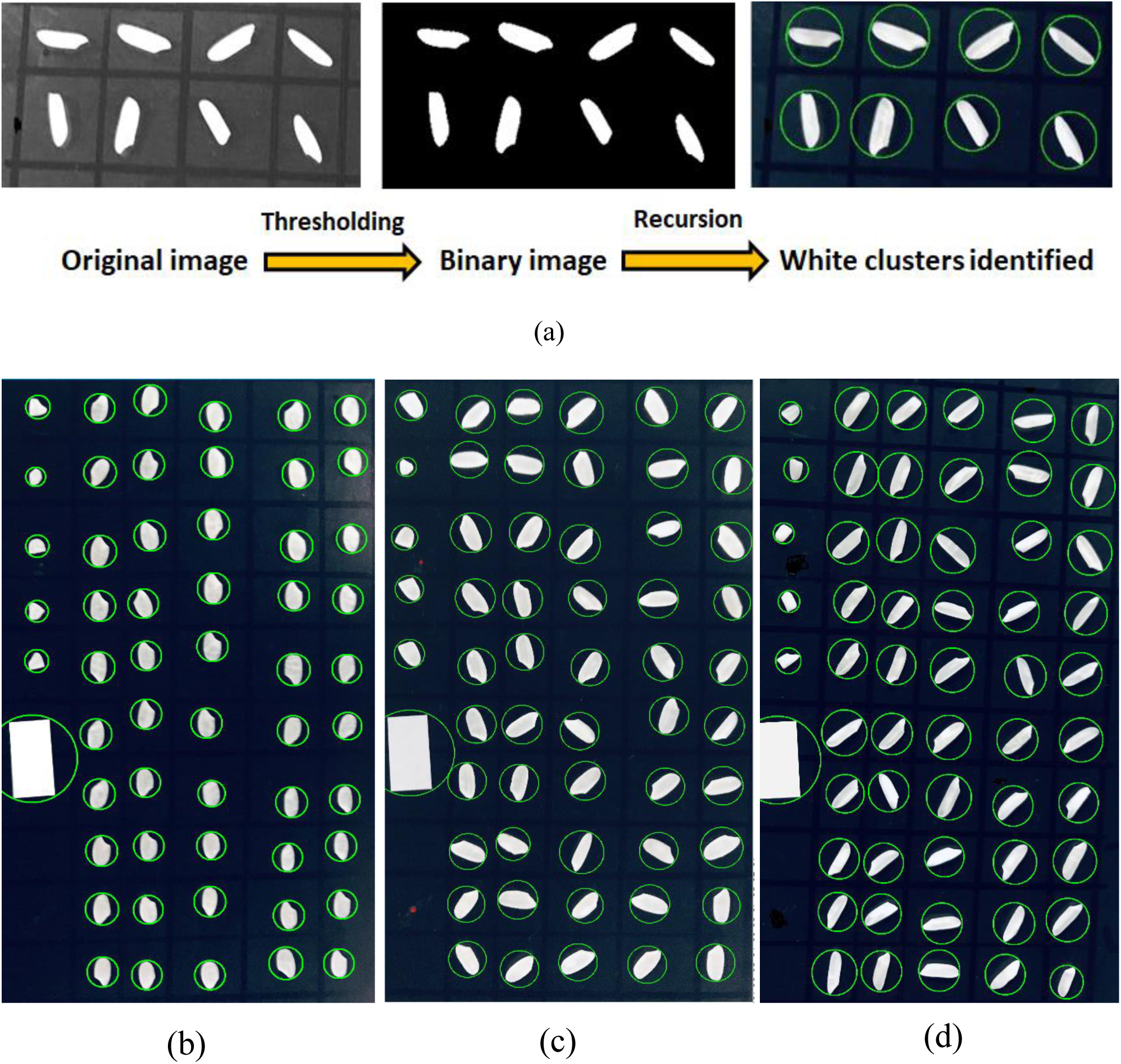
(a) Steps to identify the rice kernels in the image. The original image is converted in to binary image using a thresholding operation followed by recursive function calls to identify cluster of connected white pixel (rice kernels). Processed images for three cultivars (b) Calhikari-202, (c) Jupiter, and (d) CL153

A white paper strip (13 × 7 mm) included in every image, enabled us to calibrate and convert the image from pixel domain to real world dimensions. The size of the white strip is intentionally kept larger than any rice kernel. This allowed the code to easily identify the white strip as the largest cluster of connected white-pixels and use it to calibrate the rice kernels. In order to calculate the size of a kernel, the distance between every pixel from other pixels in the cluster was calculated. The two pixels with the largest distance were assumed to be the pixels at the both ends along the length of the kernel. The distance between these two pixels is the length of the kernel 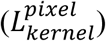 in units of pixels. A green circle of diameter equal to the length of the pixel is drawn, with its center being the midpoint between the two end pixels. Then finally the length of the kernel 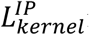 is converted from pixel to mm using Eq 3.

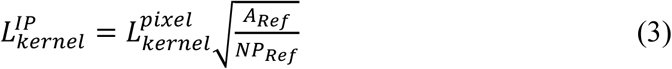

### 2.4 Machine learning

#### 2.4.1 Stacked Ensemble Model (SEM)

In order to improve the estimations of the size and mass of the individual rice kernel from the image processing, we have used a novel ML method, where an ensemble of different ML models is used to enrich the accuracy of the predictions (Singh et al., 2019). This process of pooling various ML models in layers, as the data flow forward from the input layer to the output is known as stacking and the model is called a Stacked ensemble model (SEM). A SEM typically consists of one or more layers as shown in Fig 3. A data point essentially consists of an input feature vector (independent variables) and the output properties corresponding to the input feature set. The input feature vector is fed in to the ML models of the first layer. Every ML model in the first layer makes the prediction of the output variable based on the input feature vector. The individual prediction made by each ML model in the first layer is then pooled together in the form of a *N_1_* sized vector (*V_1_*), where *N_1_* is the number of ML models in the first layer. The feature vector *V_1_* is then used as input for every ML models in the second layer. The ML models in the second layer make the predictions for the output (size and mass) of the kernel based on the predictions made by the *N_1_* ML models. This step improves with the accuracy of the predictions made by the first layer. For our SEM, we have used only two layers, the second layer being the output layer. Some of the most popular ML models, such as Artificial Neural Network (ANN), Random Forest (RF), Support Vector Machine (SVM), Kernel Ridge Regression (KRR), and k-Nearest Neighbors (k-NN) were used to construct the SEM for this study. We have used standard ML libraries of the Scikit-Learn for python.

**Fig. 3.**
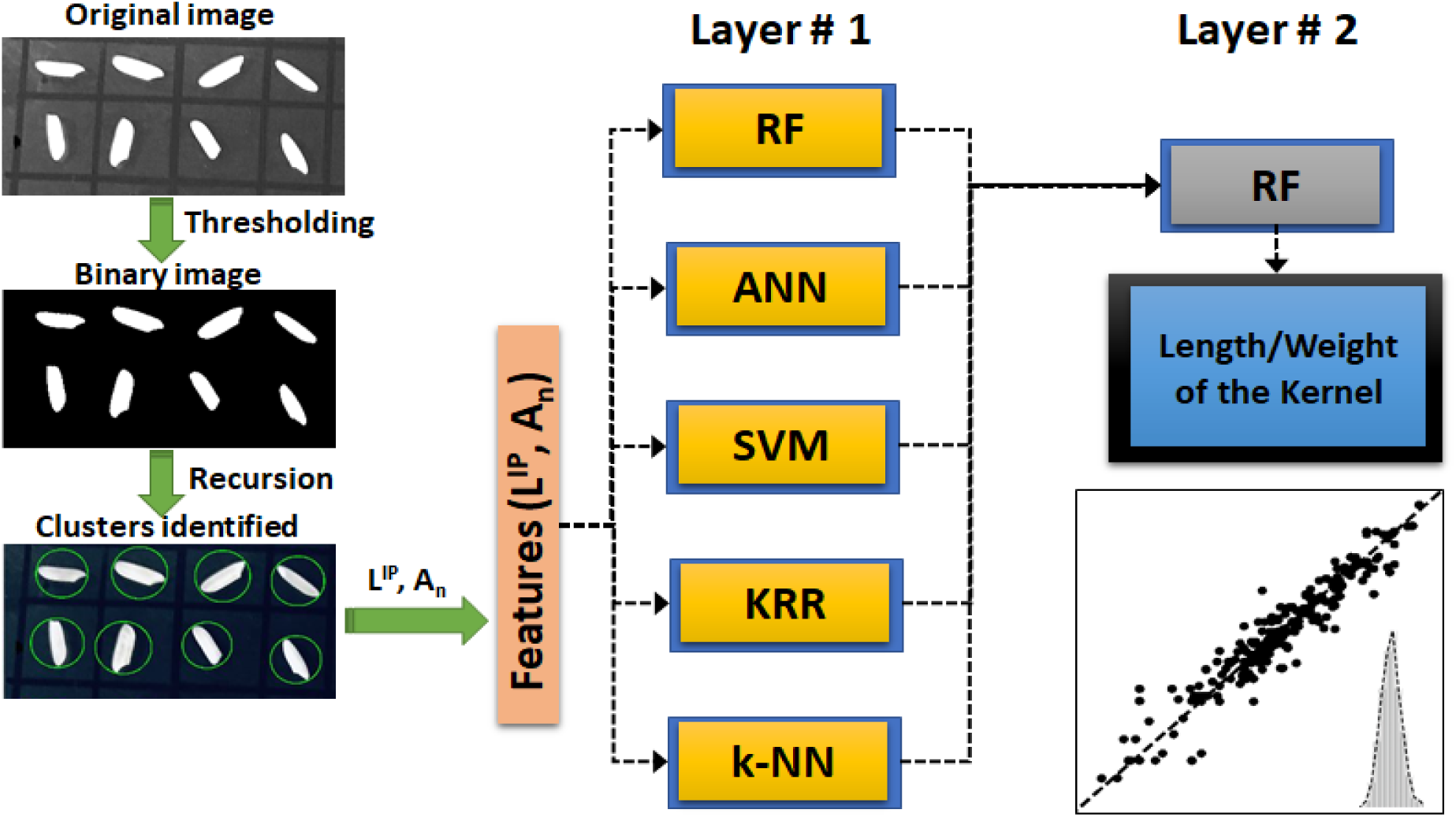
Process flow to predict the size and weight of the rice kernels. The original image is processed to create binary (black and white) image. The recursive algorithm is used to identify and count the clusters of white pixels, each cluster representing a rice kernel. The features (length and area of the cluster) are then fed in to the SEM to estimate the length and weight of the individual rice kernels in the image.

#### 2.4.2 Artificial Neural Network (ANN)

In recent years, artificial Neural Network (ANN) has been increasingly utilized in the field of science and engineering to build predictive models. Several researchers have used ANN to classify the rice grains (Fayyazi et al., 2017; Gujjar and Siddappa, 2013; Mousavi Rad et al., 2012b; Prajapati and Patel, 2013). As shown in Fig 4, an ANN model consists of several nodes and layers of nodes. A node is the compute unit, which receives input signals from the nodes of its preceding layer. The input signal values are multiplied with weighting factors and summed up.

**Fig. 4.**
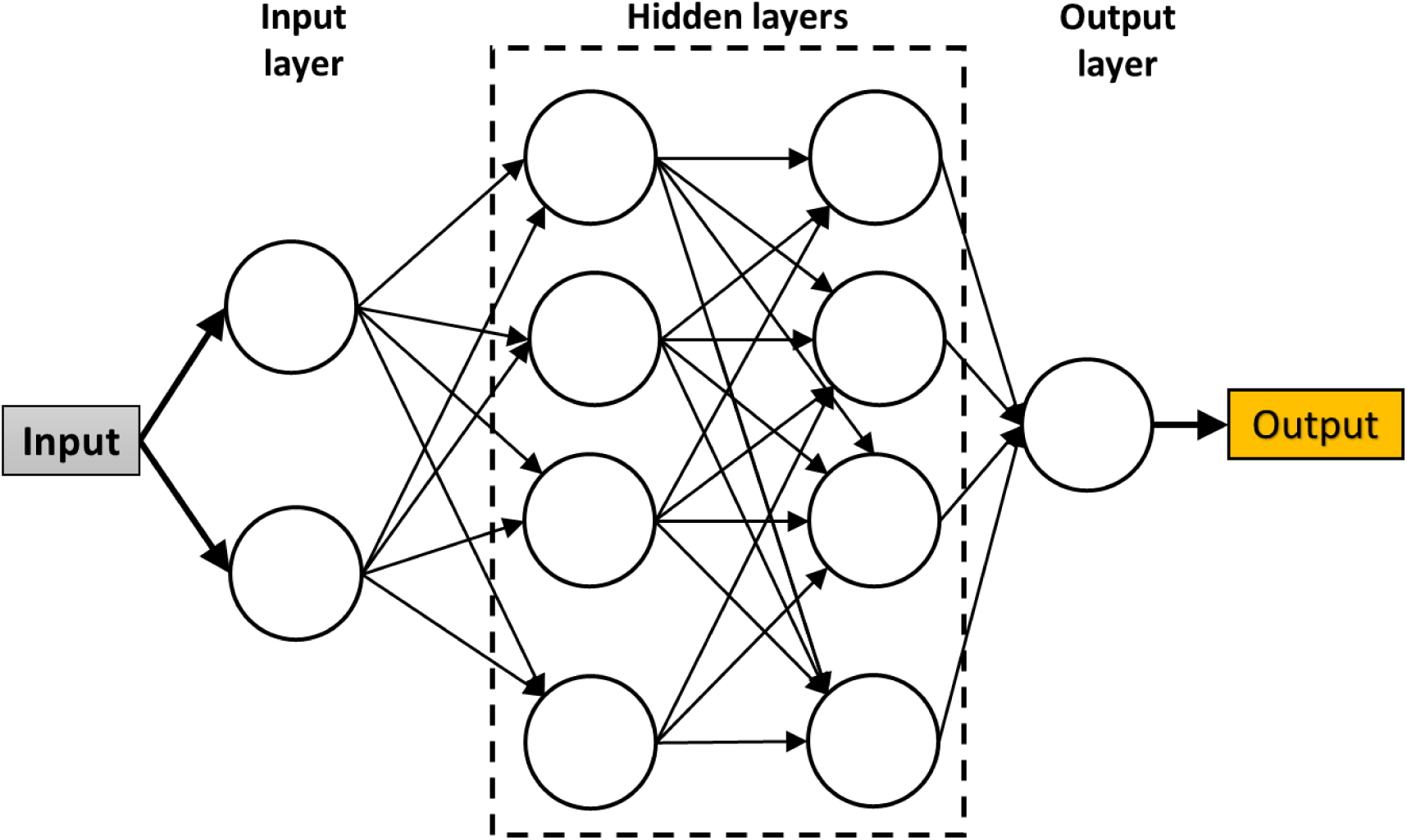
Schematic of an example artificial neural network (ANN) with two inputs, one output and two hidden layers made of 4 nodes each.

The summed signal is then passed through an activation function, which acts like a filter and modifies the signal. There are several activation functions which differ from each other based on the threshold and shape of the filter function. Some of the most popular activation functions are ReLu, ELU, sigmoid, tanh and linear (Singh and Abbassi, 2018). For this study, we have used ReLu activation function. The single modified signal from the activation function is then passed on to every node of the next layer. During the training operation, the weighting factors for each node is tuned using an optimizer to minimize the prediction error.

#### 2.4.3 Random Forest (RF)

Random forest is a popular and successful ensemble supervised ML model made of several decision tree (DT) models as shown in Fig. 5 (Quinlan, 1987; Yu et al., 2010; Silins et al., 2012; Farid et al., 2014). The RF model makes the final decision based on the voting or averaging of the underlying DT models. A DT is a logical model where the training data points are split and grouped based on their similarity. Every node in a DT consists of a logical expression which is used to decide whether a data point is sent to the adjacent node located in the left or right side of the given node. The first node of the decision tree is called *root* and the subsequent splits are called *branches*. The end nodes where data do not exist any further are called *leaves*. In order to make a prediction with a trained DT, the input data are supplied at the root node, from where the data flow through the branches based on the logical expressions until it reaches a leaf node. The output of a DT is an average of linear regression of the training data points corresponding to the leaf node. The shape and parameters of a DT are highly dependent on the training data set. The original training data are bootstrapped to generate multiple data set for training the individual DT models of the RF. The bootstrapping is a process of randomly picking data points from the original data set, one point at a time such that a data point may be picked once or more or not picked at all. In a bootstrapped data set, some points are repeated or missing from the original data set. The different data set generated from the bootstrapping steps ensure that the forest is made of diverse decision trees. The simple architecture of the underlying DT makes the RF one of the fastest ML models.

**Fig. 5.**
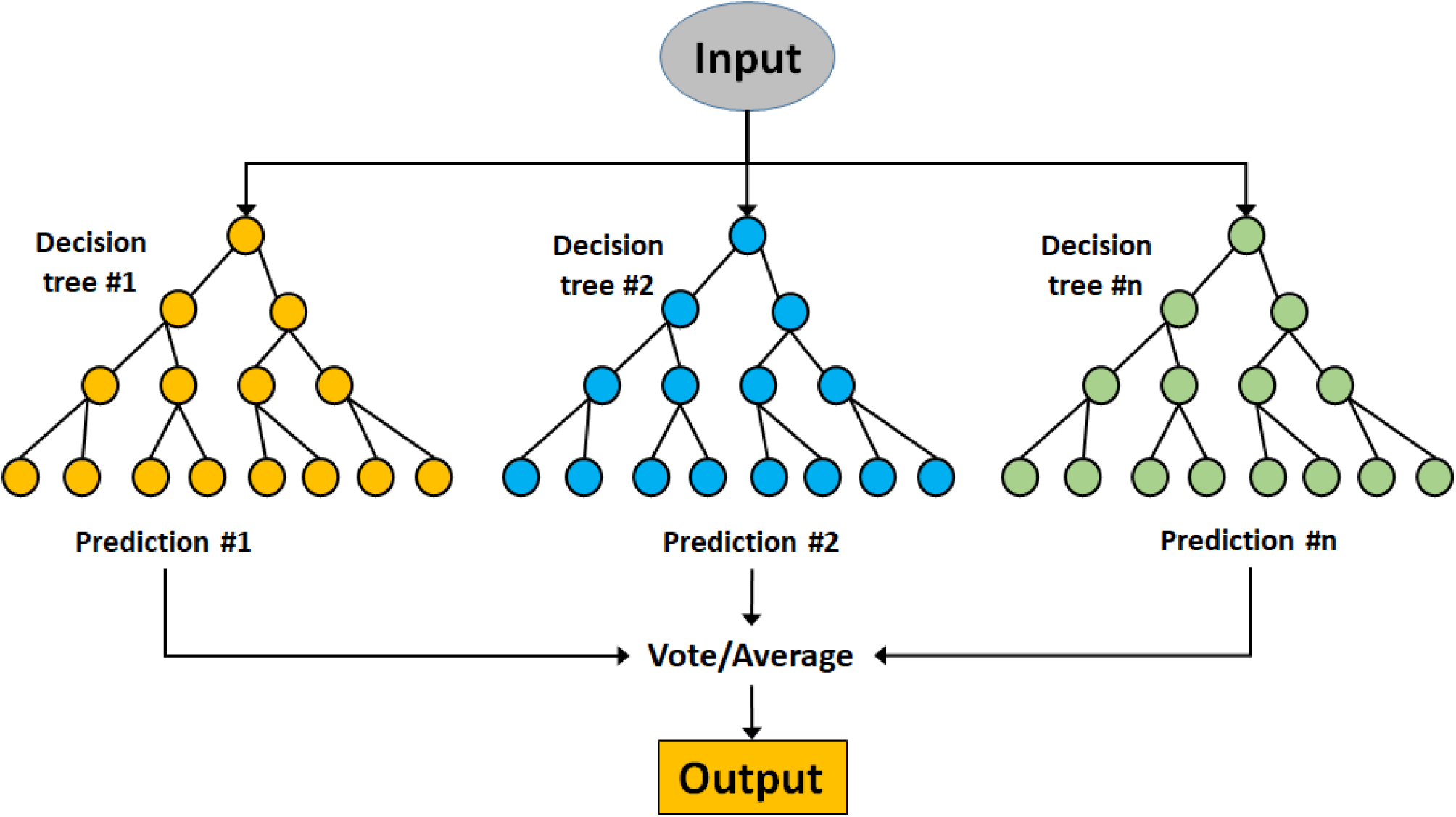
The schematic sample of a Random Forest made of n decision trees. The final decision is made by either voting or averaging the predictions from all trees in the forest.

#### 2.4.4 Support Vector Regressor (SVR)

A SVR is a regressor variant of the popular Support Vector Machine (SVM) ML model, which is typically used in classification application (Christianini and Taylor, 2000). Chen et al. (2012) and Kaur and Singh (2013) applied the SVM to classify the head rice and broken rice. The training data set may not be easy to be segregated in n-dimensional feature space using a linear hyperplane, where ‘n’ is number of features in the training data set (Fig 6a). The underlying algorithm of the SVR model transforms the input features from original n-dimensional space (F) to a higher dimensional space (M) such that the data set can be split into two or more homogeneous fragments using linear hyperplanes. During the training process, the coefficients (*a_1_, a_2_, a_3_, …, a_n_* and *b*) of the hyperplanes (Eq. 4) are optimized. The goal of the optimization is to maximize the margin (δ) between the training data points and the hyperplanes.

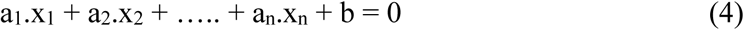

**Fig. 6.**
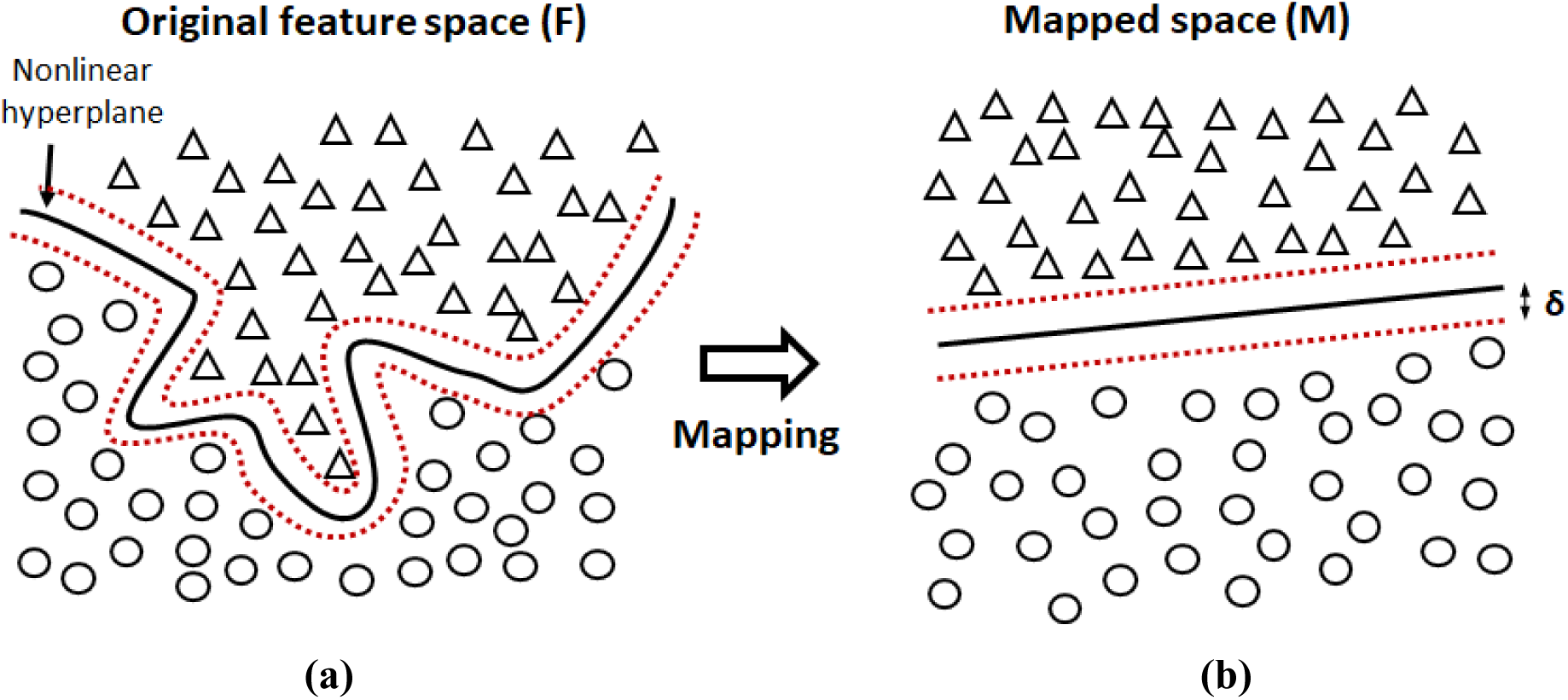
The figure shows an example where (a) data points from the original feature space F is (b) mapped onto the higher dimensional space M where the data points are separable with linear hyperplane. The solid black line represents an example hyperplane in 2D feature space and the red dotted lines represent the margin between the hyperplane and the training data points at the separation boundary.

The kernel-trick method is used to transform the training data set from original n-dimensional feature space F to a higher dimensional space M. In the transformed space M, the data points are linearly separable using the hyperplanes (Fig. 6b). For this study, we used the most popular Radial Basis Function (RBF) as described by Eq 5, where *k(x_i_,x_j_)* is the kernel function between two points *x_i_* and *x_j_* and *σ* represents the kernel width. The value of *k*(*x_i_*, *x_j_*) represents the similarity between two rice grains for our case study. For two similar rice grains, the value of *k*(*x_i_*, *x_j_*) → 1.

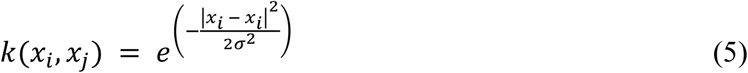

#### 2.4.5 Kernel Ridge Regression (KRR)

A Kernel Ridge Regression, similar to SVR, transforms the training data set from original feature space F to a higher dimensional space M with the help of kernel-trick method. In the higher dimensional space M, the training data are highly separable with linear hyperplanes. For example, the predicted length for rice kernel L(*x_i_*) with feature *x_i_*, can be represented by the Eq 6.

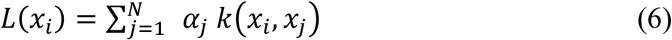

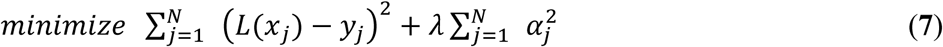

The parameter vector α is obtained by minimizing the objective function in Eq 7, where λ is a regularization penalty term and *y_i_* is the measured length of the rice kernel. The main difference between the SVR and KRR is the penalty term λ multiplied to the norm of α in the objective function to be minimized (Aurlien, 2017). This prevents the model from overfitting. KRR is not highly scalable and the speed suffers with increasing size of training data set.

#### 2.4.6 k-Nearest Neighbors (kNN)

k-nearest neighbors ML method, as the name suggests, makes the prediction based on average from the k nearest training data points to the input feature vector (Aurlien, 2017). For our study, we used three neighboring points (k=3) to make the final prediction (Fig 7). We used the Euclidean distance (Eq 8) to locate the three closest neighbors. For a given input feature vector (*A_n_* and 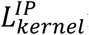), the algorithm calculates its Euclidean distance from all the training data points in 2-dimensional domain. The mass of the rice kernel is predicted by averaging the masses of the three rice kernels closely matching its *A_n_* and 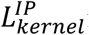 from the training data set. This method, due to its simplicity, is one of the fastest ML methods and performs well for evenly spread large data set in a featured space, especially in the region of enquiry. The downside to this method is that it stores all the training data points, which can become an issue with large training data set.

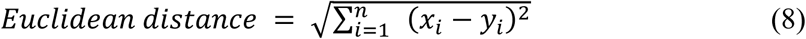

**Fig. 7.**
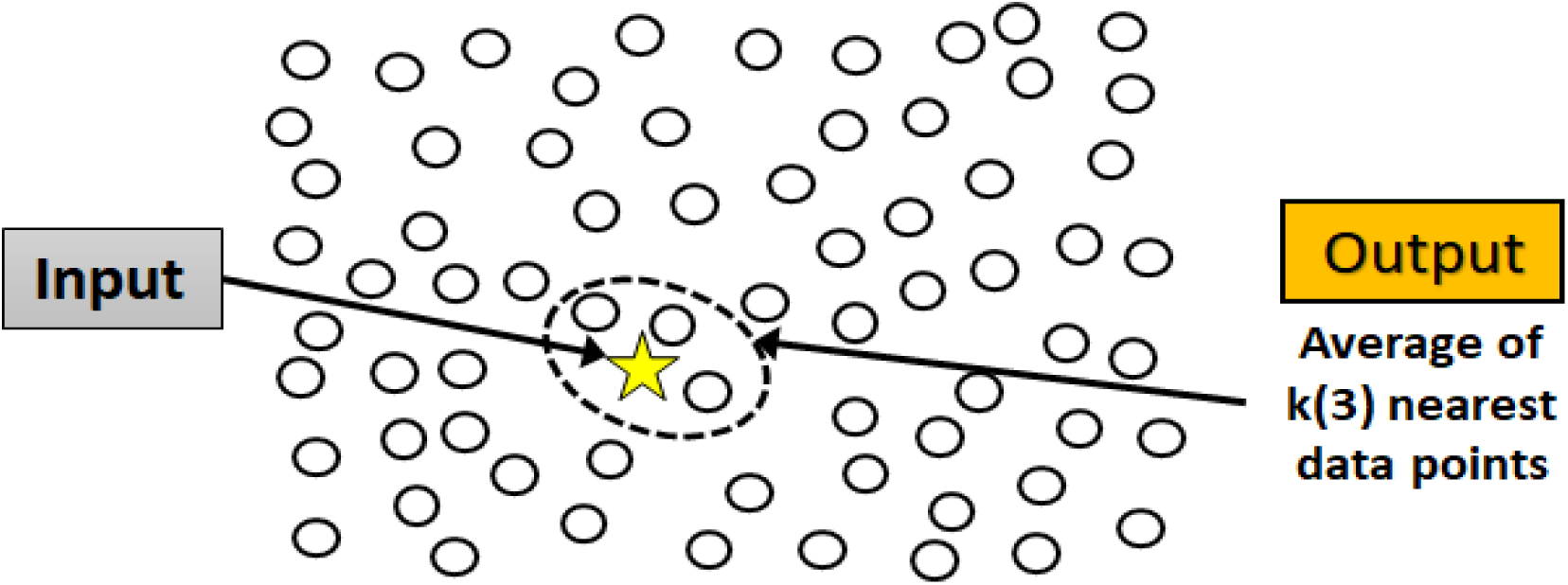
k-NN

### 2.5 Performance of SEM

The performance of any regression model is typically quantified with Root Mean Square Error (*RMSE*), Mean Absolute Percentage Error (*MAPE*) and correlation coefficient (*R^2^*) as shown in Eq 9-12. The actual value and predicted quantity are represented by ‘y’ and ‘*x’* for ‘n’ number of predictions.

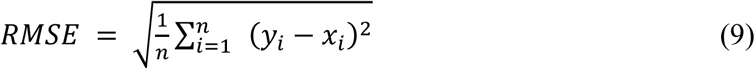

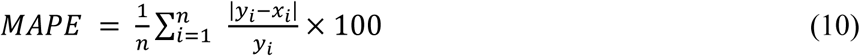

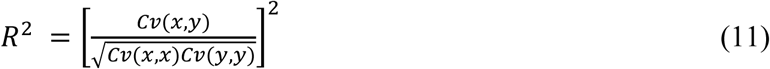

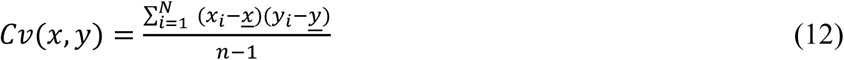

### 2.6 Uncertainty quantification of ML model (Bootstrapped statistics)

Quantification of the residual errors of a ML model is typically performed using single score indicator, such as RMSE and MAPE and it is used to understand the expected error for a prediction. These models however do not reflect the robustness of the ML model or the uncertainty associated with the predictions. Here we have used bootstrapping technique to generate several data sets and evaluate the average residual for each data set. The bootstrapping method basically takes the original data set *D_O_* of size *n* and creates *N_BS_* = 1000 new data sets, each of size n, by randomly picking data points from the original data set. A bootstrap data set can have same points from the original data set *D_O_* picked multiple times or not picked at all. The SEM was used to predict for each of 1000 bootstrapped data set. The variance of the mean residuals for each data set was used to calculate the uncertainty range. The average of the means of residuals of bootstrapped data set represents the accuracy of the ML model, whereas the variance of the mean residuals represents the robustness of the ML model. A ML model which shows narrow distribution and lower average of the mean residuals is superior and desirable.

## 3 Results and Discussion

### 3.1 Kernel size estimation by image processing algorithm

The individual kernel lengths predicted by the image processing method 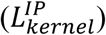 are compared with their measured values 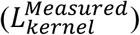 (Fig 8). The average measured kernel lengths of cultivars Calhikari, Jupiter, and CL153 were 4.66, 5.87, and 6.41 mm, respectively, whereas the corresponding sizes predicted by the image processing method were 4.54, 5.92 and 6.35 mm. We found a good correlation between the predicted and measured values with *R^2^* of 0.91, 0.94 and 0.97 for small kernel Calhikari, medium kernel Jupiter and large kernel CL153, respectively. The errors for 1000-kernel average size were 0.164 ± 0.008, 0.146 ± 0.006 and 0.150 ± 0.007 mm, respectively for the three cultivars (Fig 8). The RMSE values for the three cultivars were 0.204, 0.179 and 0.187 mm, whereas the MAPE values for those were 3.58%, 2.52% and 2.47%, respectively. The quantitative fitness indicators (RMSE, MAPE, and R^2^) for medium and long kernels (Jupiter and CL153) were marginally better than those for the smaller kernels (Calhikari).

**Fig. 8.**
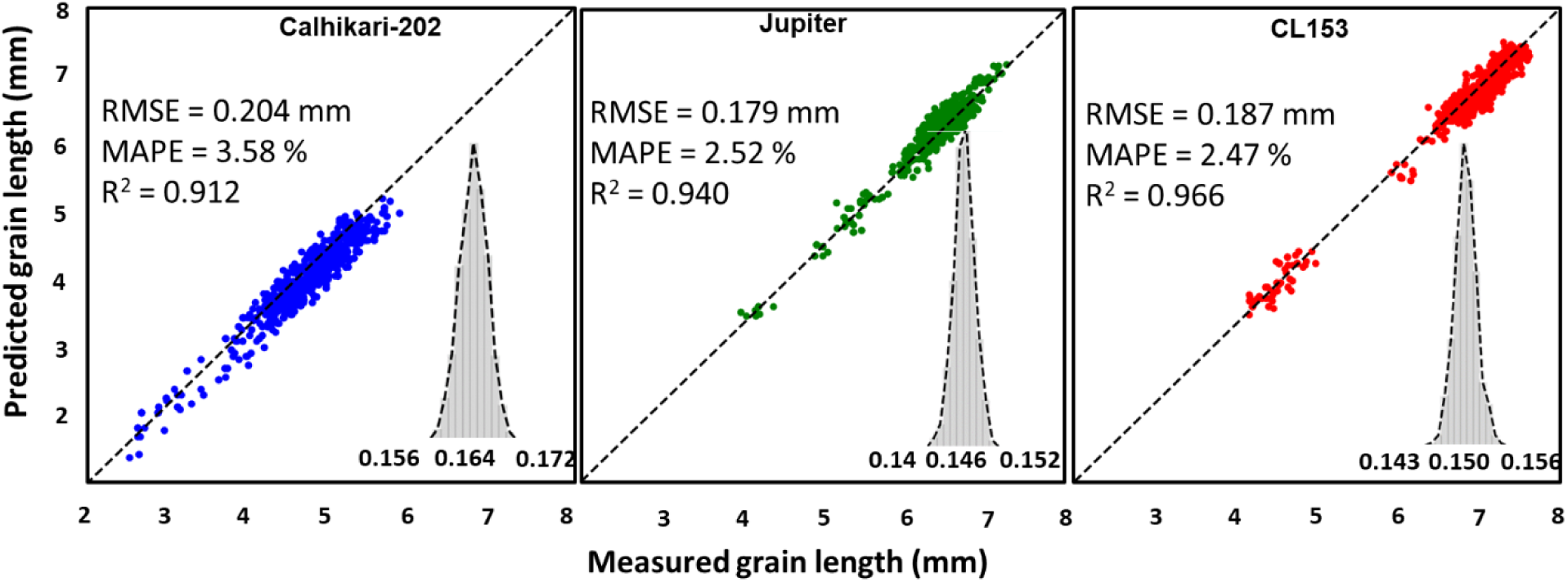
Measured kernel size vs predicted kernel size for the three rice cultivars, Calhikari-202, Jupiter, and CL153.

The differences between the 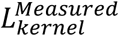 and 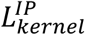 could be primarily due to two reasons, measurement errors and image processing errors. The sources of image processing errors can be further attributed to calibration error, thresholding error, and image resolution. The calibration error is a result of any error in measurements of the reference object dimensions which contributes to the errors in rice kernel size estimation. During the thresholding process, the image is converted into binary image with a threshold value. A pixel corresponding to the rice kernel may be identified as the background pixel (rgb = 0, 0, 0) and vice versa. This can lead to an under-/over-estimate of the number of pixels of a kernel. The resolution of the image plays an important role as well. An image with higher resolution has a smaller least count due to more pixels per rice kernel than the image with lower resolution. The smaller kernels are represented by fewer number of pixels than the long kernels, hence, have relatively smaller signal to noise ratio and prediction accuracy. A camera with higher image resolution could be used to improve the prediction accuracy further. In addition to the image processing errors, the measurement errors are also higher for the smaller objects.

### 3.2 Kernel size estimated by SEM

The dataset of 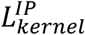 and 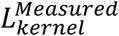 was used to train the SEM model. The SEM model was designed to take 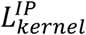 as input and predict the size of the kernel 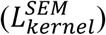 as the output. The predictions made by SEM model are marginally better than the 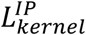 values as shown in Table 1. The prediction accuracy of the individual ML models in first layer of the SEM model are further improved by the ML model of the second layer. The SEM model performs better with the medium and long grains than the short grains. The average lengths of three cultivars as predicted by SEM are 4.69 mm, 5.81 mm and 6.37 mm, respectively for Calhikari-202, Jupiter and CL153. The RMSE, MAPE and R^2^ values generated from SEM model for all the tested cultivars are shown in fig 9-11, which reveal that the SEM model further improves the prediction results overall. For example, the RMSE value generated from predicted model for Calhikari-202 improved from 0.204 to 0.161 mm in SEM model (RF: Layer 1), which was further improved to 0.160 mm by the ML model of the second layer (RF: Layer 2). Similar trends were noticed for MAPE and R^2^. For the same cultivar, MAPE improved from 3.58% to 2.838% and R^2^ from 0.912 to 0.932 in SEM model with the further improvement to 2.819% and 0.934, respectively in the second layer. These trends were common for the other two cultivars, however, the improvements in the prediction indicators were superior in cultivars Jupiter and CL153 compared to Calhikari-202. The RMSE, MAPE and R^2^ of software and SEM predicted models for the grain sizes of tested rice cultivars are shown in Table 1. The uncertainty quantification of residuals (mean error ± 95% limit) also showed progressive improvements in the accuracy with the layers of ML models in the SEM model. For example, the mean residual error for cultivar CL153 improved from 0.150 ± 0.007 mm to 0.14 ± 0.01 mm in SEM model (RF: Layer 1), which further improved to 0.13 ± 0.01 mm in the second layer (RF: Layer 2). Similar trend was noticed for the other two cultivars.

**Fig. 9.**
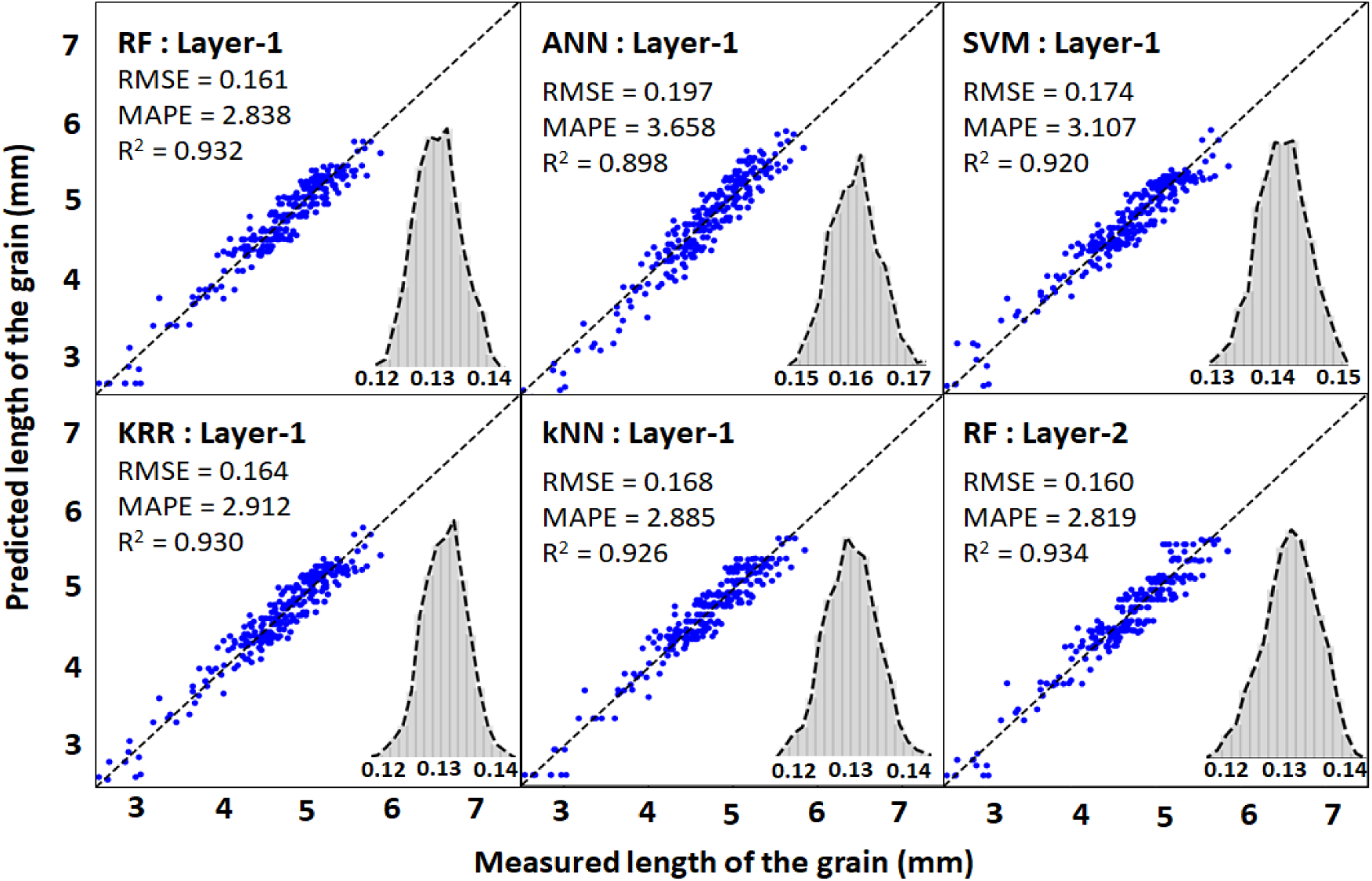
Cultivar: SEM Cahlikari-202

**Fig. 10.**
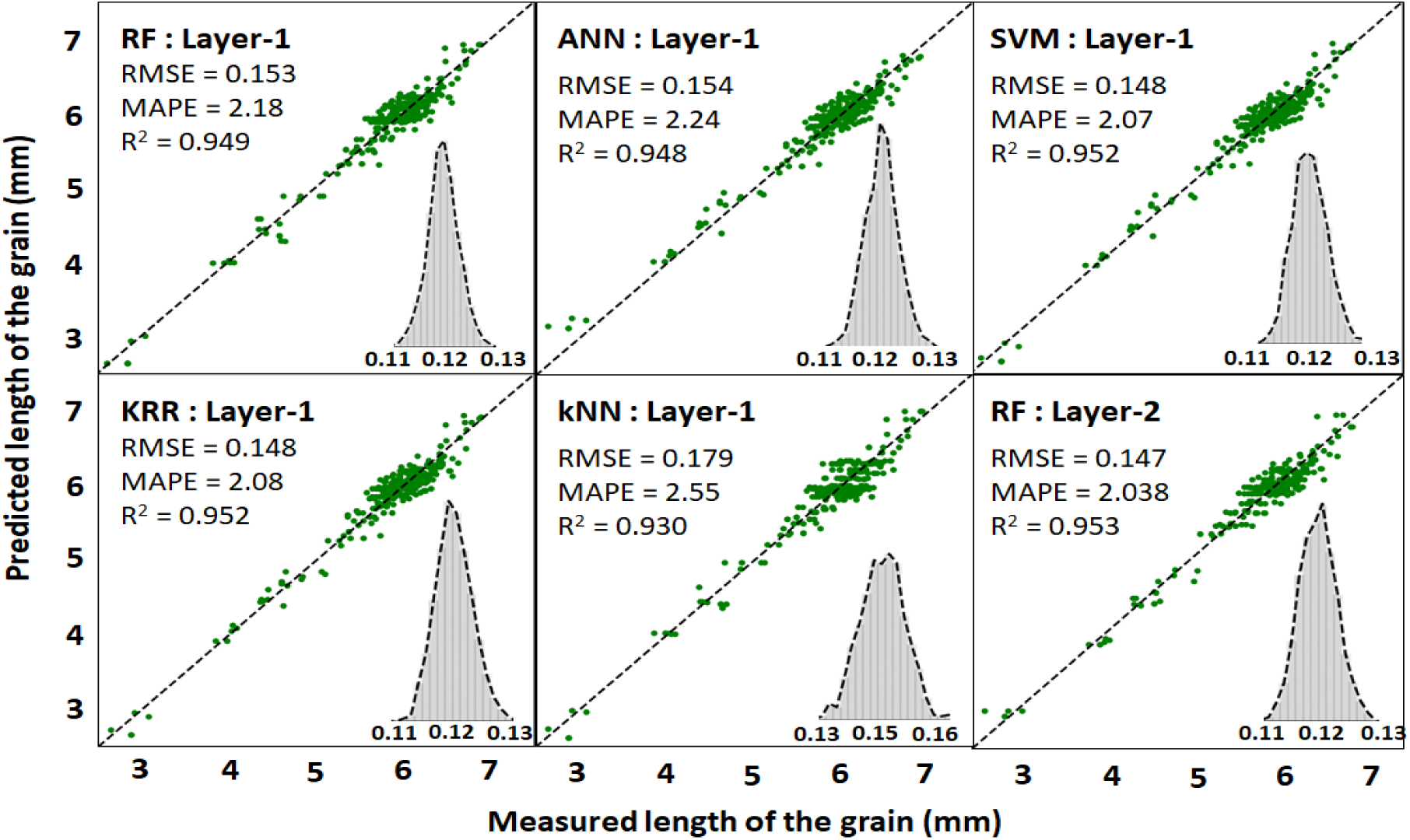
Cultivar: SEM Jupiter

**Fig. 11.**
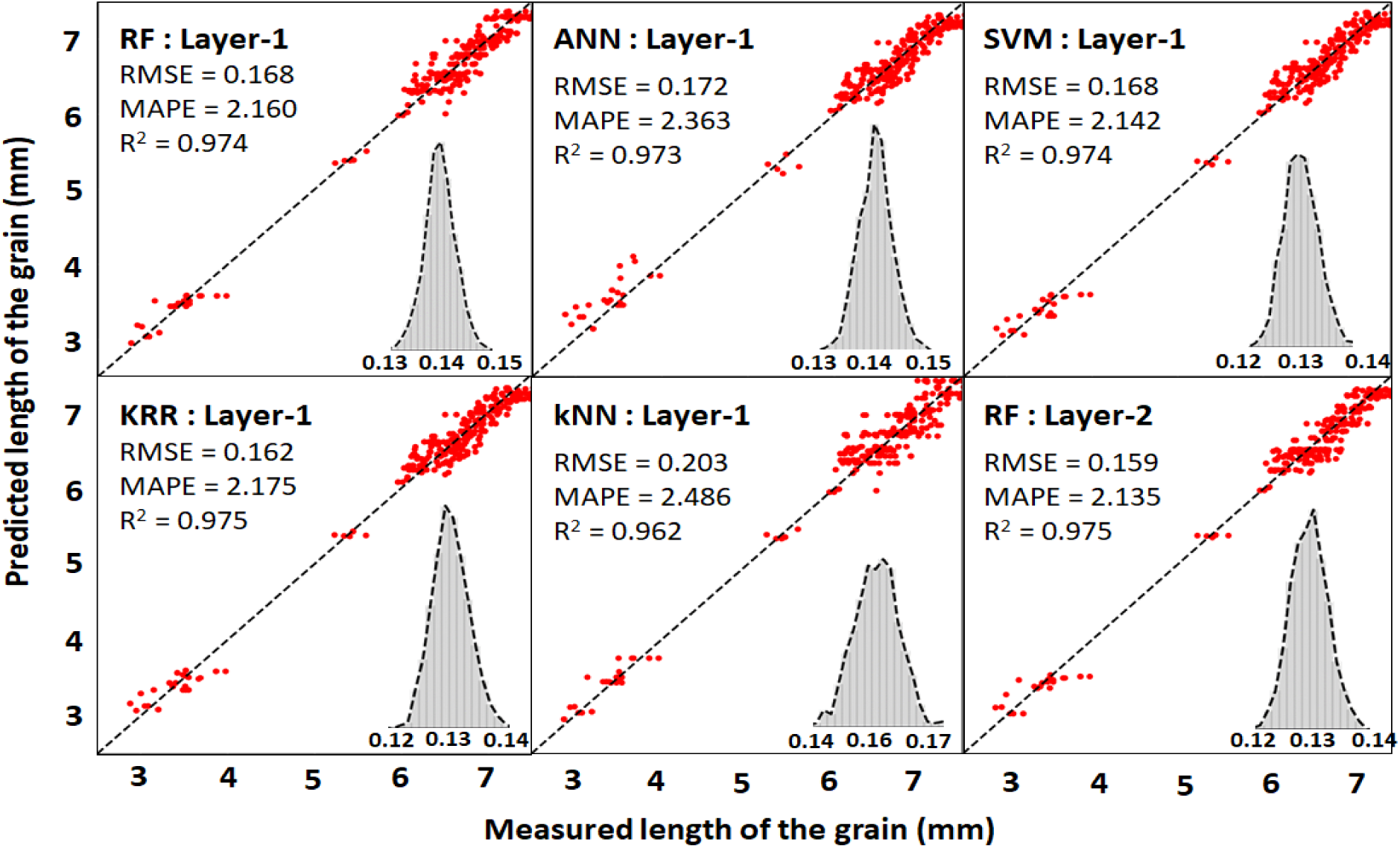
Cultivar: SEM CL153

**Table 1.** Performance indicators for software predictions and ML models for the size (mm) of the rice kernels.

| Image Processing Prediction Stats |  |  |  |  | SEM Predictions Stats |  |  |  |  |
| --- | --- | --- | --- | --- | --- | --- | --- | --- | --- |
| Cultivars | RMSE (mm) | MAPE (%) | R<sup>2</sup> | Uncertainty quantification of resifuals (Mean error ± 95% limit) |  | RMS E (mm) | MAPE (%) | R<sup>2</sup> | Uncertainty quantification of residuals (Mean error ± 95% limit) |
| Calhikari-202 | 0.204 | 3.58 | 0.912 | 0.164 ± 0.008 | Layer 1: RF | 0.161 | 2.838 | 0.932 | 0.13 ± 0.01 |
|  |  |  |  |  | Layer 1: ANN | 0.197 | 3.658 | 0.898 | 0.16 ± 0.01 |
|  |  |  |  |  | Layer 1: SVM | 0.174 | 3.107 | 0.92 | 0.14 ± 0.01 |
|  |  |  |  |  | Layer 1: KRR | 0.164 | 2.912 | 0.93 | 0.13 ± 0.01 |
|  |  |  |  |  | Layer 1: k-NN | 0.168 | 2.885 | 0.926 | 0.13 ± 0.01 |
|  |  |  |  |  | Layer 2: RF | 0.16 | 2.819 | 0.934 | 0.13 ± 0.01 |
| Jupiter | 0.179 | 2.52 | 0.94 | 0.146 ± 0.006 | Layer 1: RF | 0.153 | 2.18 | 0.949 | 0.12 ± 0.01 |
|  |  |  |  |  | Layer 1: ANN | 0.154 | 2.24 | 0.948 | 0.12 ± 0.01 |
|  |  |  |  |  | Layer 1: SVM | 0.148 | 2.07 | 0.952 | 0.12 ± 0.01 |
|  |  |  |  |  | Layer 1: KRR | 0.148 | 2.08 | 0.952 | 0.12 ± 0.01 |
|  |  |  |  |  | Layer 1: k-NN | 0.179 | 2.55 | 0.93 | 0.15 ± 0.01 |
|  |  |  |  |  | Layer 2: RF | 0.147 | 2.038 | 0.953 | 0.12 ± 0.01 |
| CL153 | 0.187 | 2.47 | 0.966 | 0.150 ± 0.007 | Layer 1: RF | 0.168 | 2.16 | 0.974 | 0.14 ± 0.01 |
|  |  |  |  |  | Layer 1: ANN | 0.172 | 2.363 | 0.973 | 0.14 ± 0.01 |
|  |  |  |  |  | Layer 1: SVM | 0.168 | 2.142 | 0.974 | 0.13 ± 0.01 |
|  |  |  |  |  | Layer 1: KRR | 0.162 | 2.175 | 0.975 | 0.13 ± 0.01 |
|  |  |  |  |  | Layer 1: k-NN | 0.203 | 2.486 | 0.962 | 0.16 ± 0.01 |
|  |  |  |  |  | Layer 2: RF | 0.159 | 2.135 | 0.975 | 0.13 ± 0.01 |

### 3.3 Kernel weight estimation by image processing algorithm

We hypothesize that the 2D projected image of a rice kernel, as exposed to the camera, represents the two major dimensions of the kernel and can be used as a variable to predict the mass of the kernel. The pixel area (number of pixels) representing a kernel is standardized by dividing it with the number of pixels of the reference object of known dimensions (the rectangular paper strip). The standardized pixel area is then normalized by multiplying it with a constant *C_n_* = 4.5 so that the largest kernel in our measurement sample has a normalized pixel area (*A_n_*) close to 1.0 (Eq 11).

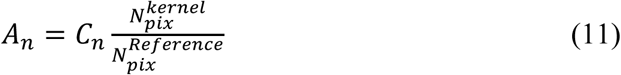

The measured mass of the individual kernel was correlated with its *A_n_*. The average measured kernel weights of cultivars Calhikari, Jupiter and CL153 were 16.82 mg, 19.81 mg and 17.13 mg and their respective normalized pixel area were 0.543, 0.738 and 0.627, respectively. The *R^2^* values (>0.9) indicate a good correlation between the *A_n_* and the measured weight of the rice grains for all three cultivars as shown in Table 2 and fig 12. The *R^2^* values for small kernel Calhikari, medium kernel Jupiter and large kernel CL153 were 0.908, 0.951 and 0.922, respectively, which show a stronger correlation for longer grains (Jupiter and CL153) than the shorter grain (Calhikari-202), indicating a slightly lower prediction accuracy for shorter grains.

**Fig. 12.**
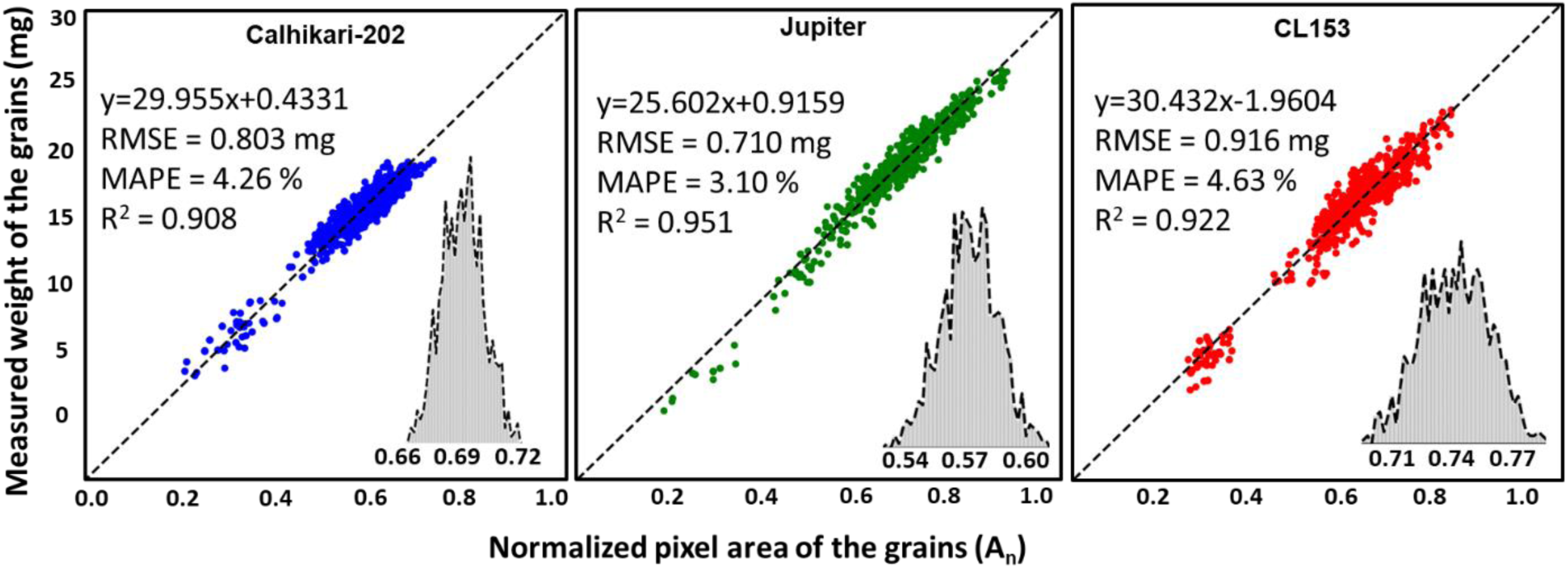
Normalized pixel area (*A_n_*) of rice kernels vs measured weight of kernels

**Table 2.** Performance indicators for image processing predictions and ML models for the mass of the rice kernels.

| Image Processing Prediction Stats |  |  |  |  | SEM Predictions Stats |  |  |  |  |
| --- | --- | --- | --- | --- | --- | --- | --- | --- | --- |
| Cultivars | RMSE (mg) | MAPE (%) | R<sup>2</sup> | Uncertainty quantification of residuals (Mean error ± 95% limit) |  | RMSE (mg) | MAPE (%) | R<sup>2</sup> | Uncertainty quantification of residuals (Mean error ± 95% limit) |
| Calhikari-202 | 0.803 | 4.26 | 0.908 | 0.69 ± 0.03 | Layer 1: RF | 0.695 | 3.287 | 0.932 | 0.51 ± 0.05 |
|  |  |  |  |  | Layer 1: ANN | 0.772 | 4.11 | 0.916 | 0.64 ± 0.04 |
|  |  |  |  |  | Layer 1: SVM | 0.791 | 4.16 | 0.912 | 0.66 ± 0.05 |
|  |  |  |  |  | Layer 1: KRR | 0.776 | 4.124 | 0.915 | 0.65 ± 0.04 |
|  |  |  |  |  | Layer 1: k-NN | 0.931 | 4.348 | 0.878 | 0.67 ± 0.07 |
|  |  |  |  |  | Layer 2: RF | 0.596 | 2.554 | 0.95 | 0.40 ± 0.05 |
| Jupiter | 0.71 | 3.1 | 0.951 | 0.57 ± 0.03 | Layer 1: RF | 0.525 | 2.03 | 0.973 | 0.37 ± 0.04 |
|  |  |  |  |  | Layer 1: ANN | 0.713 | 2.896 | 0.95 | 0.53 ± 0.05 |
|  |  |  |  |  | Layer 1: SVM | 0.688 | 2.83 | 0.954 | 0.51 ± 0.05 |
|  |  |  |  |  | Layer 1: KRR | 0.651 | 2.72 | 0.958 | 0.50 ± 0.05 |
|  |  |  |  |  | Layer 1: k-NN | 0.837 | 3 | 0.931 | 0.55 ± 0.07 |
|  |  |  |  |  | Layer 2: RF | 0.471 | 1.65 | 0.978 | 0.31 ± 0.04 |
| CL153 | 0.916 | 4.63 | 0.922 | 0.74 ± 0.03 | Layer 1: RF | 0.816 | 3.61 | 0.954 | 0.54 ± 0.06 |
|  |  |  |  |  | Layer 1: ANN | 0.868 | 4.323 | 0.947 | 0.60 ± 0.06 |
|  |  |  |  |  | Layer 1: SVM | 0.849 | 4.301 | 0.95 | 0.66 ± 0.06 |
|  |  |  |  |  | Layer 1: KRR | 0.857 | 4.393 | 0.949 | 0.68 ± 0.05 |
|  |  |  |  |  | Layer 1: k-NN | 0.804 | 3.442 | 0.955 | 0.55 ± 0.06 |
|  |  |  |  |  | Layer 2: RF | 0.728 | 3.08 | 0.963 | 0.48 ± 0.06 |

The errors for 1000-kernel average weight were 0.69 ± 0.03, 0.57 ± 0.03 and 0.74 ± 0.03 mg, respectively for the three cultivars (fig 12). The RMSE values for the three cultivars were 0.803 g, 0.71 mg and 0.916 mg, whereas the MAPE values for those were 4.26%, 3.1% and 4.63%, respectively. The quantitative fitness indicators (RMSE, MAPE and R^2^) reflect an unusual trend, indicating a better fitness model for medium size grain (Jupiter) than the long grains (CL153). The higher errors and lower prediction accuracy in the smaller kernels could be attributed to the smaller number of pixels overall in the smaller kernels than the longer kernels. In general, rice kernels were wider in one of its two radial directions, of which, the smaller breadth was ignored in this study.

The 1000 grains average weights of three cultivars as predicted by SEM are 16.76, 20.12, and 17.25 mg, respectively for Calhikari-202, Jupiter, and CL153. The SEM model improved the estimations for the masses of the kernels compared to the linear least-square fit model. The prediction accuracy indicators (RMSE, MAPE and R^2^) obtained from SEM models for the kernel mass of all the cultivars are shown in Table 2 and fig 13-15, which clearly show the progressive improvement of the accuracy with the layers of ML models in the SEM model. For example, the RMSE value generated from predicted model for Jupiter reduced from 0.71 mg to 0.525 mg in SEM model (RF: Layer 1), which was further decreased to 0.471 mg by the ML model of the second layer (RF: Layer 2). Similar trends were noticed for MAPE and R^2^. For the same cultivar, MAPE improved from 3.1% to 2.03% and R^2^ from 0.951 to 0.973 in SEM model with the further improvement to 1.65 and 0.978, respectively, in the second layer. These trends were common for the other two cultivars. The uncertainty quantification of residuals (mean error ± 95% limit) also showed progressive improvements in the accuracy with the layers of ML models in the SEM model. For example, the uncertainty quantification of residuals for cultivar Calhikari-202 improved from 0.69 ± 0.003 to 0.51 ± 0.05 mg in SEM model (RF: Layer 1), which further improved to 0.40 ± 0.05 mg in the second layer (RF: Layer 2). Similar trend was noticed for the other two cultivars.

**Fig. 13.**
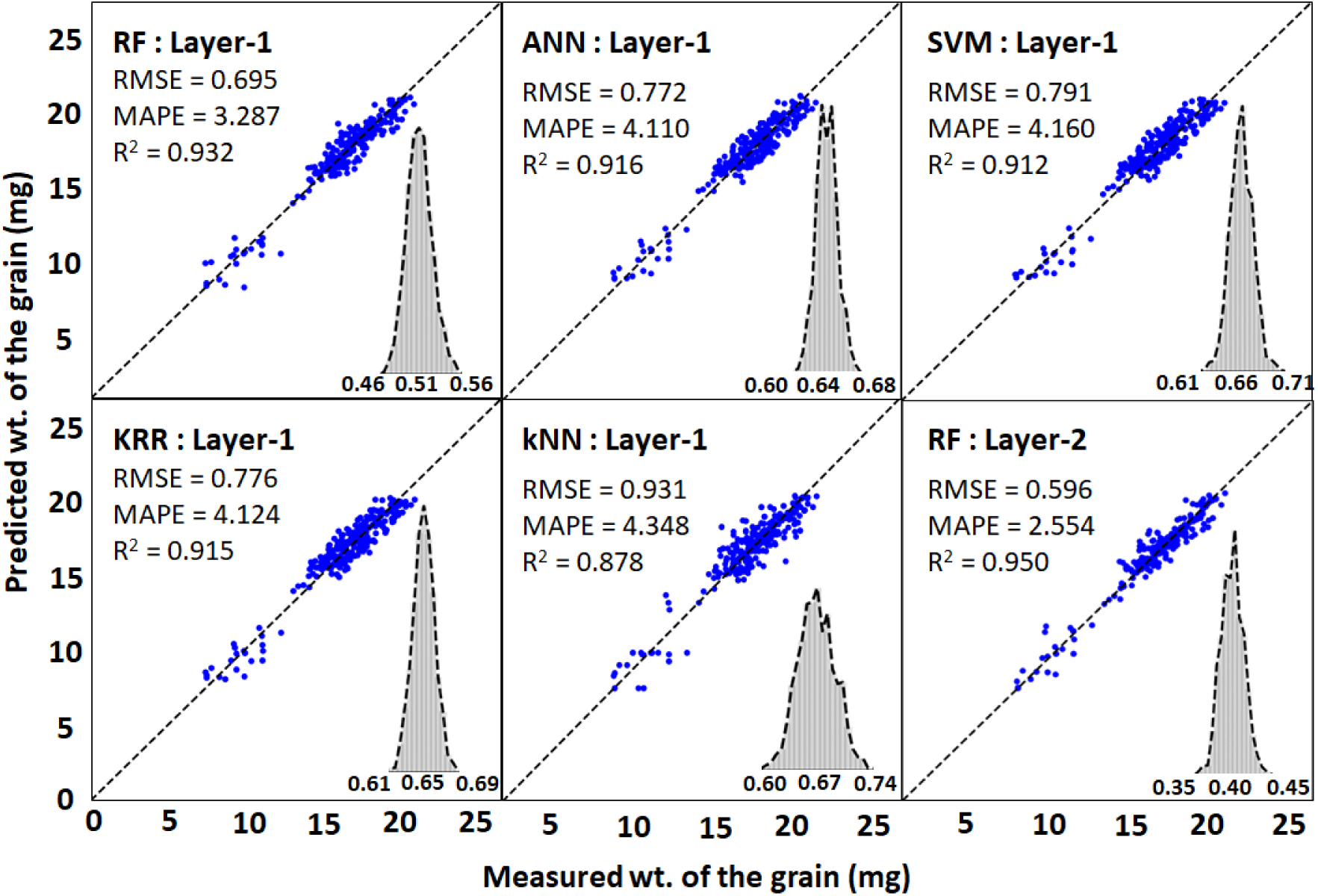
SEM Calhikari-202

**Fig. 14.**
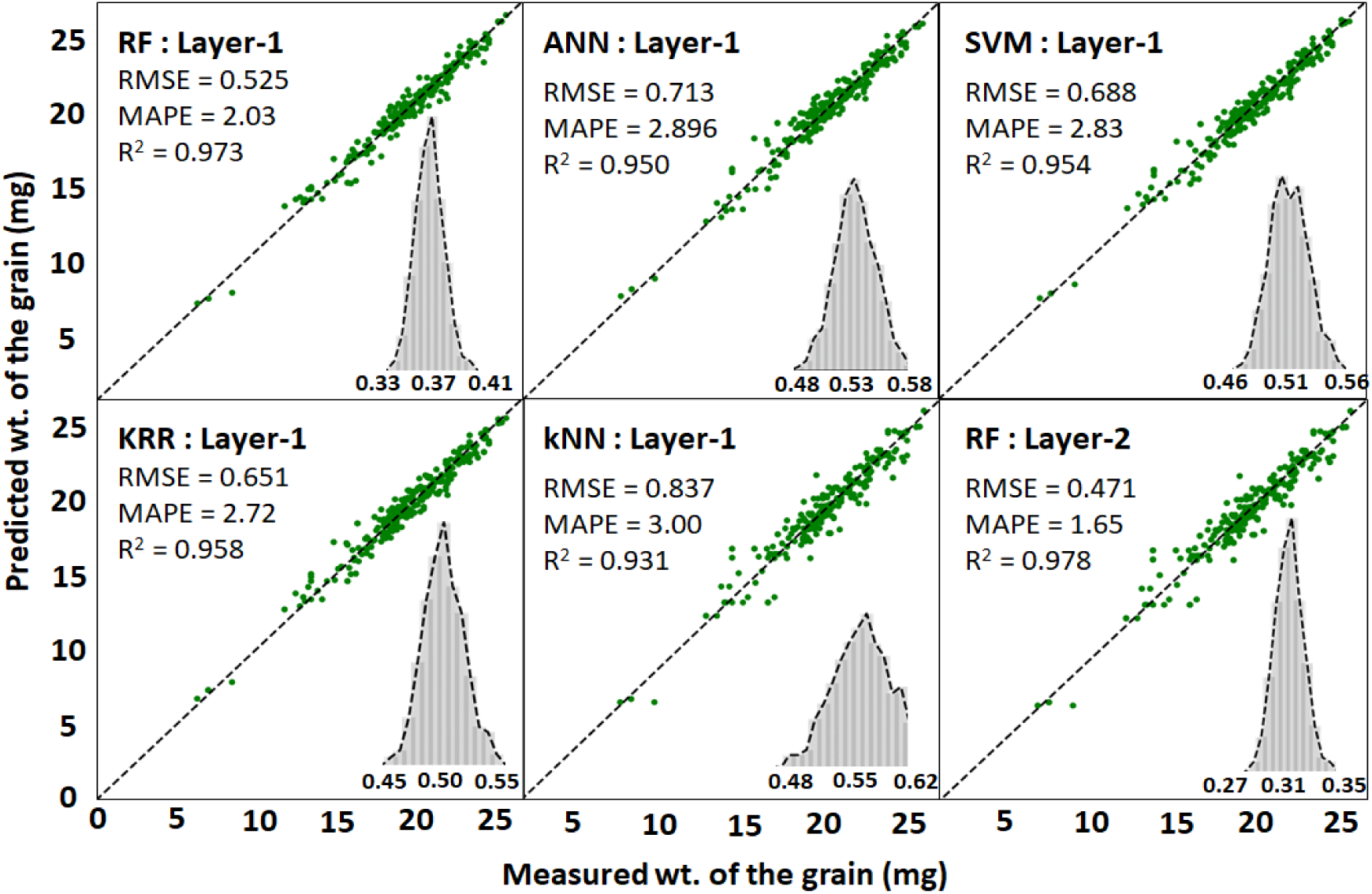
Cultivar: SEM Jupiter

**Fig. 15.**
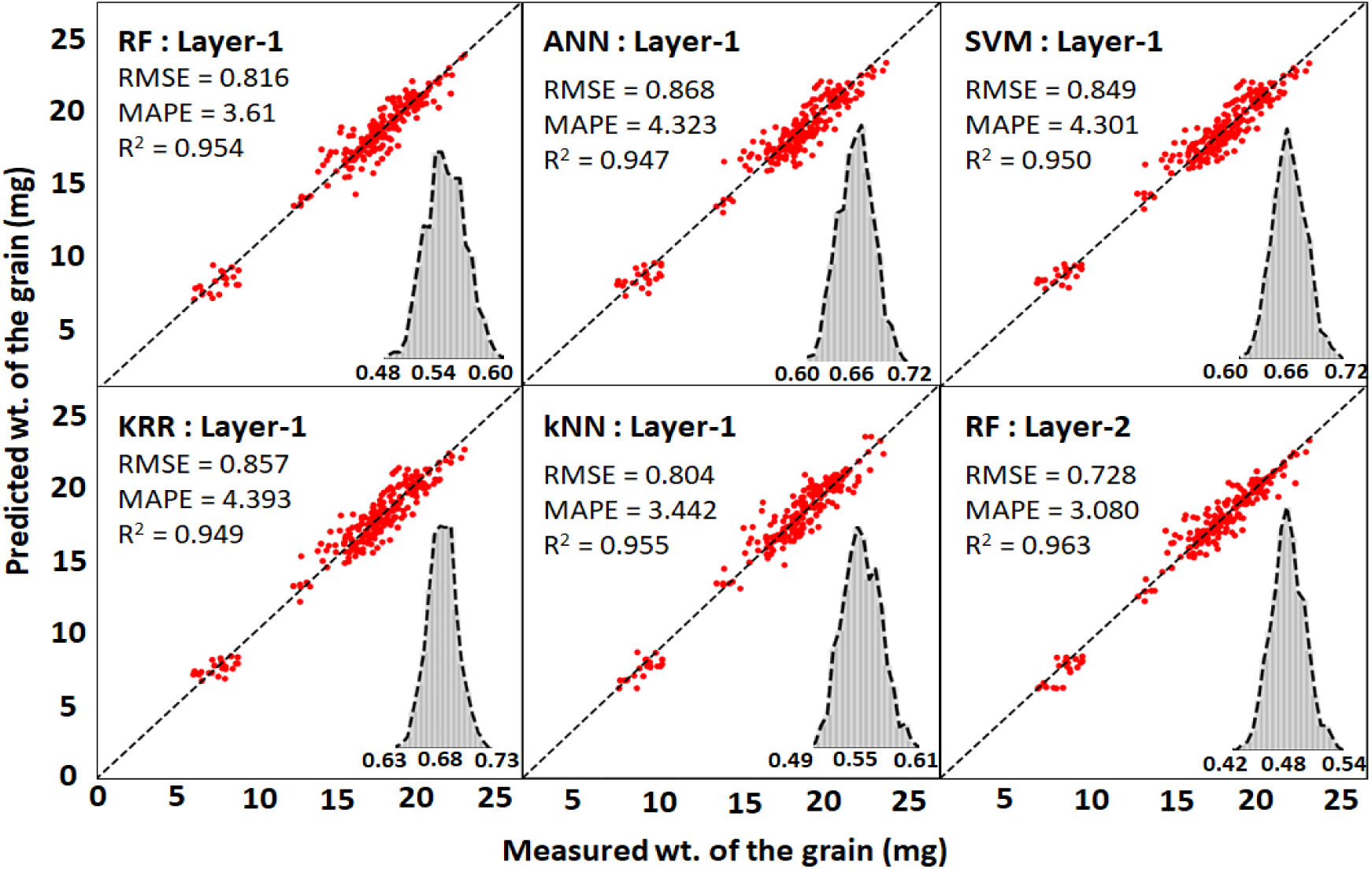
Cultivar: SEM CL153

Since, the breadths (in both dimensions) of the rice kernels could be an important factor contributing to the weight-area prediction correlation, ignoring the smaller breadth could be a possible reason for unusual trend in RSME and MAPE values for the cultivars. The differences in RMSE, MAPE and *R^2^* for the cultivars could primarily be attributed to the measurement, calibration, and thresholding errors, as well as image resolution. In addition, the error in the measurements of the reference dimensions also contributes to the errors in rice kernel size estimation, which could ultimately reflect in the *A_n_* and then propagate to correlation between *A_n_* and their corresponding measured weights. As stated in the previous section, during the thresholding process, a pixel belonging to the rice kernel may be identified as the background pixel (rgb=0,0,0), which can lead to an underestimation of the number of pixels corresponding to a rice kernel. Similarly, a few background pixels around the rice grain could be identified as white pixels and be included in the cluster of pixels for the rice, leading to an overestimation of the *A_n_*.

In addition, the resolution of the image too contributes to the overall error. With higher number of pixels, a high-resolution image has a smaller value of the least count and hence a higher accuracy in capturing the size and shape of the kernel. The higher resolution image also captures the fine details of the shape with higher accuracy. An image with lower resolution describes the image of kernels with fewer coarser pixels which can either underestimate or overestimate the value of 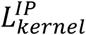 and *A_n_*. In order to explore the effect of image resolution, we studied a test case with one rice kernel and compared its 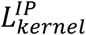 and *A_n_* values at different image resolutions as shown in Fig 16. The coarsening operation of the original image was performed with an image processing tool GIMP, using a linear scaling method. The relative difference (RD) between quantities calculated using the lower resolution images and the original image (100% resolution) converged at around 75% resolution. The number of pixels representing the rice kernel in the original image is 11,391 pixels and for 75% resolution image is 6,792 pixels. We can conclude that the image processing error due to resolution is minimal if each full kernel in an image is represented by approximately 7,000 pixels or greater. The percentage image resolutions are calculated with respect to the original image of the rice kernel (Eq 12-13). The Fig 16 demonstrates that our image setting was sufficient to minimize the error due to image resolution.

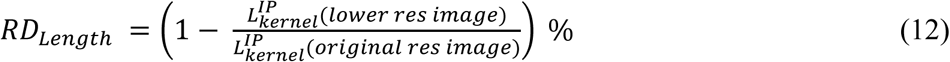

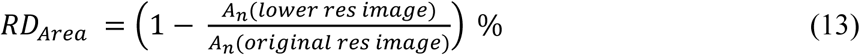

**Fig. 16.**
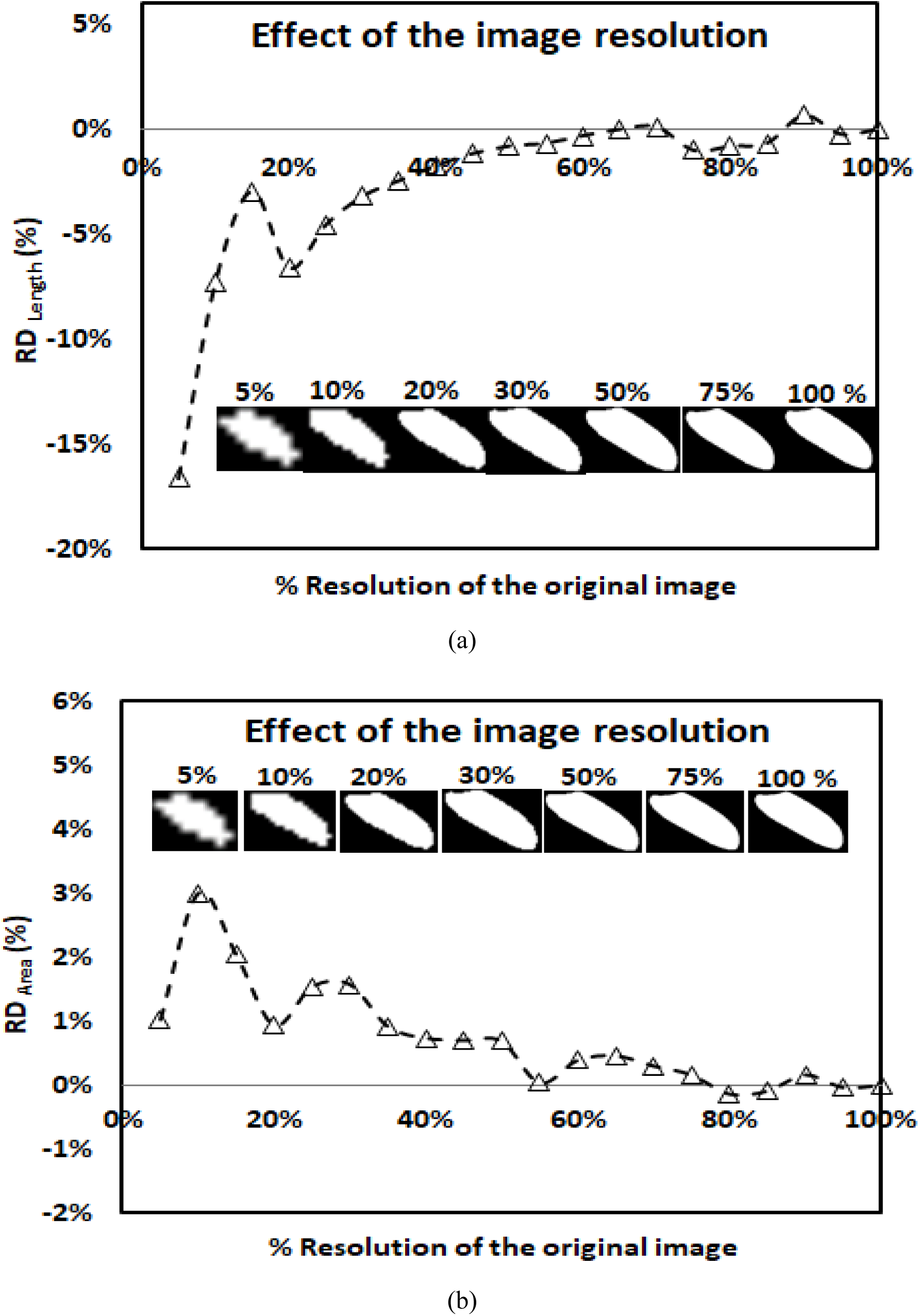
Relative difference due to the effect of image resolution on the calculated values of (a) 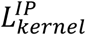 and (b) *A_n_* value for a rice kernel. The insat images show the same rice kernel at different % resolutions of the original image.

## 4 Conclusions

We have successfully demonstrated a novel application of image processing and ML to accurately measure the size and mass of individual rice kernels with high confidence level. Through our image processing algorithm, we were able to obtain the size and area of the individual rice kernels in the form of pixels. The pixel information was then converted into feature vectors, each feature vector representing the size and area of an individual kernel. The pixels corresponding to each rice kernel is the key parameter and serves as the basis for predicting the length and weight of the kernels. We stacked several ML models to create a SEM, which could enrich the prediction accuracy from one layer to another. The final prediction accuracy of the SEM was superior to any individual member ML model or least square error fit, demonstrating the power of stacking the ML models. The effect of image resolution on the image processing error was also studied. We found that the errors due to resolution converged for a rice kernel which was represented by greater than approximately 7,000 pixels.

We used three rice cultivars namely Calhikari-202, Jupiter, and CL153 to evaluate our methodology. The average measured length of the 1000 kernels were 4.66, 5.87 and 6.41 mm, respectively. Using our image processing algorithm, we were able to estimate the average length of the 1000 kernels to be 4.54 mm, 5.92 mm, and 6.35 mm and bootstrapped residuals with 95% confidence interval of 0.16 ± 0.01 mm, 0.15 ± 0.01 mm, and 0.15 ± 0.01 mm for Calhikari-202, Jupiter and CL153 rice cultivars, respectively. Using the SEM, we were able to further improve our predictions of average lengths to be 4.69 mm, 5.81 mm, and 6.37 mm and bootstrapped residual of 0.12 ± 0.01 mm, 0.12 ± 0.01, and 0.13 ± 0.01 mm, respectively. The numbers clearly indicate that we are able to predict the length of the kernels with very high accuracy.

The average mass of the individual kernels for the three varieties were estimated to be 16.82, 19.81 and 17.03 g using the SEM. The bootstrapped residuals with 95% confidence level. For the three varieties were 0.40 ± 0.05, 0.31 ± 0.04 and 0.48 ± 0.06 mg. The low values of mean errors and narrow ranges of the 95% confidence intervals indicate the robustness and accuracy of our novel methodology. This methodology can have important applications in the field of rice processing, replacing the manual and tedious appraisal methods typically associated with kernel sample measurements, saving enormous amount of time and effort. Additionally, the use of the developed method in image analyzers could provide an objective assessment of head rice yield of bulk rice samples.

## Acknowledgement

We would like to express our gratitude to Dr Bhagwati Prakash for his valuable suggestions throughout this research.

